# Novel mechanistic targets of Forkhead box Q1 transcription factor in human breast cancer cells

**DOI:** 10.1101/2020.05.26.117176

**Authors:** Su-Hyeong Kim, Eun-Ryeong Hahm, Krishna B. Singh, Shivendra V. Singh

## Abstract

The transcription factor forkhead box Q1 (FoxQ1), which is overexpressed in different solid tumors, has emerged as a key player in the pathogenesis of breast cancer by regulating epithelial-mesenchymal transition, maintenance of cancer-stem like cells, and metastasis. However, the mechanism underlying oncogenic function of FoxQ1 is still not fully understood. In this study, we compared the RNA-seq data from FoxQ1 overexpressing SUM159 cells with that of empty vector-transfected control (EV) cells to identify novel mechanistic targets of this transcription factor. Consistent with published results in basal-like subtype, immunohistochemistry revealed upregulation of FoxQ1 protein in luminal-type human breast cancer tissue microarrays when compared to normal mammary tissues. Many previously reported transcriptional targets of FoxQ1 (e.g., *E-cadherin, N-cadherin, fibronectin 1*, etc.) were verified from the RNA-Seq analysis. FoxQ1 overexpression resulted in downregulation of genes associated with cell cycle checkpoints, M phase, and cellular response to stress/external stimuli as evidenced from the Reactome pathway analysis. Consequently, FoxQ1 overexpression resulted in S, G2M and mitotic arrest in basal-like SUM159 and HMLE cells, but not in luminal-type MCF-7 cells. There were differences in expression of cell cycle-associated proteins between FoxQ1 overexpressing SUM159 and MCF-7 cells. Finally, we show for the first time that FoxQ1 is a direct transcriptional regulator of interleukin (IL)-1α, IL-8, and vascular endothelial growth factor in breast cancer cells. Chromatin immunoprecipitation revealed FoxQ1 occupancy at the promoters of *IL-1α, IL-8*, and *VEGF*. In conclusion, the present study reports novel mechanistic targets of FoxQ1 in human breast cancer cells.

## Introduction

Breast cancer remains an alarming health concern for women worldwide reflected by more than 40,000 deaths each year in the United States alone (1). A challenging aspect in clinical management of breast cancer relates to molecular heterogeneity of the disease that is characterized by distinct gene expression signatures and overexpression of driver oncogenic proteins (2–5). Majority of the invasive mammary ductal carcinomas are broadly grouped into four subtypes, including luminal A type [estrogen receptor positive (ER+^3^), progesterone receptor positive (PR+), and human epidermal growth factor receptor 2 positive (HER2+)], luminal B type (ER+/PR+/HER2-), HER2-enriched, and basal-like (2,3). Nearly 75% of basal-like breast tumors are triple-negative due to lack of ER, PR, and HER2 expression (6). Further characterization of the disease subtype-independent oncogenic dependencies in breast cancer is necessary to identify novel druggable targets to broaden therapeutic options for different subtypes of breast cancer.

The transcription factor forkhead box Q1 (FoxQ1) has recently emerged as a key player in the pathogenesis of breast cancer (7–9). Zhang *et al*. (7) were the first to demonstrate a role for FoxQ1 in epithelial-mesenchymal transition (EMT) in breast cancer metastasis using a cross-species gene expression profiling strategy. Forced expression of FoxQ1 in a human mammary epithelial cell line (HMLE) and EpRas cells increased their ability to migrate and invade *in vitro*, and these effects were reversible by knockdown of this protein in 4T1 mouse mammary carcinoma cells (7). Moreover, FoxQ1 overexpressing EpRas cells exhibited increased propensity for pulmonary metastasis *in vivo* (7). Promotion of the EMT phenotype by FoxQ1 overexpression was due to direct repression of E-cadherin protein expression (7). In another study, FoxQ1 expression was shown to correlate with high-grade basal-like breast cancers and associated with poor clinical outcomes (8). RNA interference of FoxQ1 in a highly invasive basal-like human breast cancer cell line (MDA-MB-231) attenuated EMT phenotype and decreased its invasion ability (8). Ectopic expression of FoxQ1 in a human mammary epithelial cell line immortalized by hTERT and SV40 large T antigen (HMLER) also promoted stem-like phenotype characterized by increased mammosphere multiplicity (8). Importantly, suppression of FoxQ1 expression in MDA-MB-231 cells resulted in increased active caspase-3 level in response to treatment with several chemotherapy drugs, including 5-fluorouracil, taxol, and camptothecin (8). Transcriptional inactivation of E-cadherin by FoxQ1 overexpression was also demonstrated in this study (8). Subsequently, platelet-derived growth factor receptor α/β were identified as additional downstream targets of FoxQ1 in promotion of breast cancer stem cell-like phenotype as well as chemoresistance (9). We have also shown previously that FoxQ1 promotes breast cancer stem-like phenotype in a luminal-type (MCF-7) and a basal-like (SUM159) cell line by causing direct transcriptional repression of the tumor suppressor Dachshund homolog 1 (DACH1) (10).

Published studies thus far clearly indicate that FoxQ1 regulates expression of multiple cancer-relevant genes to promote cell invasion/migration, stem-like phenotype, and chemotherapy resistance *in vitro* and metastasis *in vivo* in breast cancer cell lines, but the possibility of additional functionally important targets of this transcription factor can’t be fully ignored. To address this question, we compared RNA-Seq data from FoxQ1 overexpressing SUM159 (hereafter abbreviated as FoxQ1 cells) cells with that of empty vector transfected control cells (hereafter abbreviated as EV cells) to identify its additional mechanistic targets. The present study reveals novel downstream targets of FoxQ1 in breast cancer including cell cycle control, interleukin (IL)-1α, IL-8, and vascular endothelial growth factor (VEGF).

## Results

### FoxQ1 protein is overexpressed in luminal-type human breast cancers compared to normal mammary tissues

We have shown previously that the level of FoxQ1 protein and mRNA is relatively higher in basal-like human breast cancer cell lines (e.g., MDA-MB-231) in comparison with a normal mammary epithelial cell line (MCF-10A) or luminal-type cells (e.g., MCF-7 and MDA-MB-361) (10). Moreover, basal-like human breast cancers exhibited higher level of FoxQ1 protein when compared to normal mammary tissues (10). Initially, we compared the expression of *FoxQ1* gene expression in white *versus* black breast cancer patients and between ductal and lobular carcinomas in breast cancer TCGA dataset. The expression of *FoxQ1* was marginally but significantly higher in the tumors of black women when compared to white women (Fig. 1*A*). On the other hand, the expression of *FoxQ1* was comparable between ductal and lobular breast cancers (Fig. 1*A*). In this study, we also examined the expression of FoxQ1 protein in tissue microarrays consisting of luminal-type breast cancers and normal mammary tissues. Fig. *1B* shows immunohistochemical staining for FoxQ1 in representative normal mammary tissues and luminal-type breast cancers. The H-score for FoxQ1 expression was higher by 1.6-fold in luminal-type breast cancers when compared to normal mammary tissues (Fig. 1*C*). These results indicated overexpression of the FoxQ1 protein in luminal-type human breast cancers.

**Figure 1.**
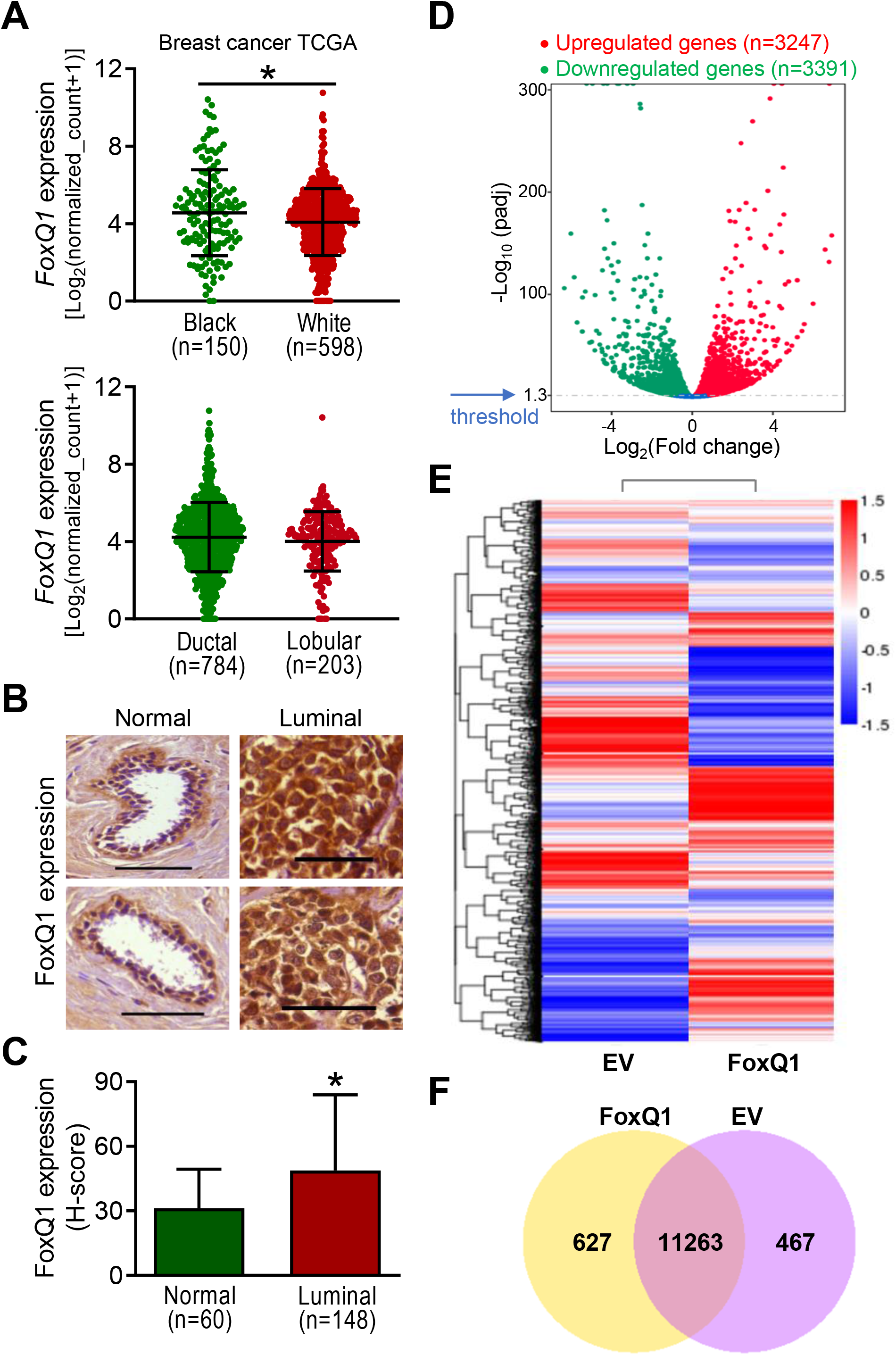
FoxQ1 expression was significantly higher in luminal-type human breast cancer specimens compared to normal human mammary tissues. ***A*,** expression of *FoxQ1* in mammary tumors of black and white women, and in ductal and lobular bacrcinomas. ***B*,** representative immunohistochemical images (200× magnification; scale bar = 100 μm) for FoxQ1 expression in normal human mammary tissues and luminal-type human breast cancer tissues. ***C*,** quantification of FoxQ1 expression in normal human mammary tissues and luminal-type human breast cancer tissues. Results shown are mean ± S.D. *P < 0.05 by two-sided Student’s t-test. ***D*,** volcano plot for differential gene expression. ***E*,** heatmap of the differentially expressed genes in empty vector transfected cells (EV) and FoxQ1 overexpressing SUM159 cells. ***F*,** venn diagram showing unique and overlapping genes between EV and FoxQ1 overexpressing SUM159 cells.

### FoxQ1-regulated transcriptome in SUM159 cells

Next, we performed RNA-Seq analysis using SUM159 cells to identify additional targets of FoxQ1. The mapping results are summarized in Table 1. The Volcano plot in Fig. *1D* depicts distribution of differentially expressed genes between FoxQ1 overexpressing SUM159 cells and EV cells with a threshold cut-off of 1.3-fold change in expression and at an adjusted *p* value of < 0.05 (Fig. 1*D*). The heatmaps of three replicates of each group exhibiting highly consistent transcriptional changes are shown in Fig. *1E*. The Venn diagram shown in Fig. 1*F* shows unique and overlapping gene expression between FoxQ1 overexpressing SUM159 cells and EV cells.

**Table 1.** The summary of mapping result

| Sample name | EV-SUM159 |  | FoxQ1-SUM159 |  |
| --- | --- | --- | --- | --- |
|  | Mean | SD | Mean | SD |
| Total reads | 65198263 | 4386610 | 72027681 | 10214885 |
| Total mapped | 58914439<br>(90.38%) | 3655237<br>(0.51%) | 64895915<br>(90.17%) | 8672753<br>(0.82%) |
| Multiple mapped | 1147142<br>(1.76%) | 81213<br>(0.01%) | 1224173<br>(1.70%)* | 154648<br>(0.03%)* |
| Uniquely mapped | 57767297<br>(88.62%) | 3574643<br>(0.52%) | 63671741<br>(88.46%) | 8518408<br>(0.80%) |
| Reads map to '+' | 28883648<br>(44.31%) | 1787322<br>(0.26%) | 31835871<br>(44.23%) | 4259204<br>(0.40%) |
| Reads map to '-' | 28883648<br>(44.31%) | 1787322<br>(0.26%) | 31835871<br>(44.23%) | 4259204<br>(0.40%) |
| Non-splice reads | 32843918<br>(50.39%) | 2036952<br>(0.30%) | 36497716<br>(50.68%) | 5119388<br>(0.56%) |
| Splice reads | 24923378<br>(38.24%) | 1546835<br>(0.33%) | 27174025<br>(37.79%) | 3464790<br>(1.14%) |

\*, Statistically significant analyzed by unpaired t test ( $P < 0.05$ ).

### Kyoto encyclopedia of genes and genomes (KEGG) pathway analysis

The top 5 pathways with upregulated genes from the KEGG analysis included pathways in cancer (84 genes), mitogen-activated protein kinase signaling (56 genes), regulation of actin cytoskeleton (49 genes), focal adhesion (50 genes), and axon guidance (37 genes) (Fig. 2*A*). The top 5 pathways with downregulated genes from the KEGG pathway analysis were cell cycle (41 genes), oocyte meiosis (38 genes), progesterone-mediated oocyte maturation (29 genes), lysine degradation (18 genes), and protein processing in endoplasmic reticulum (46 genes) (Fig. 2*B*).

**Figure 2.**
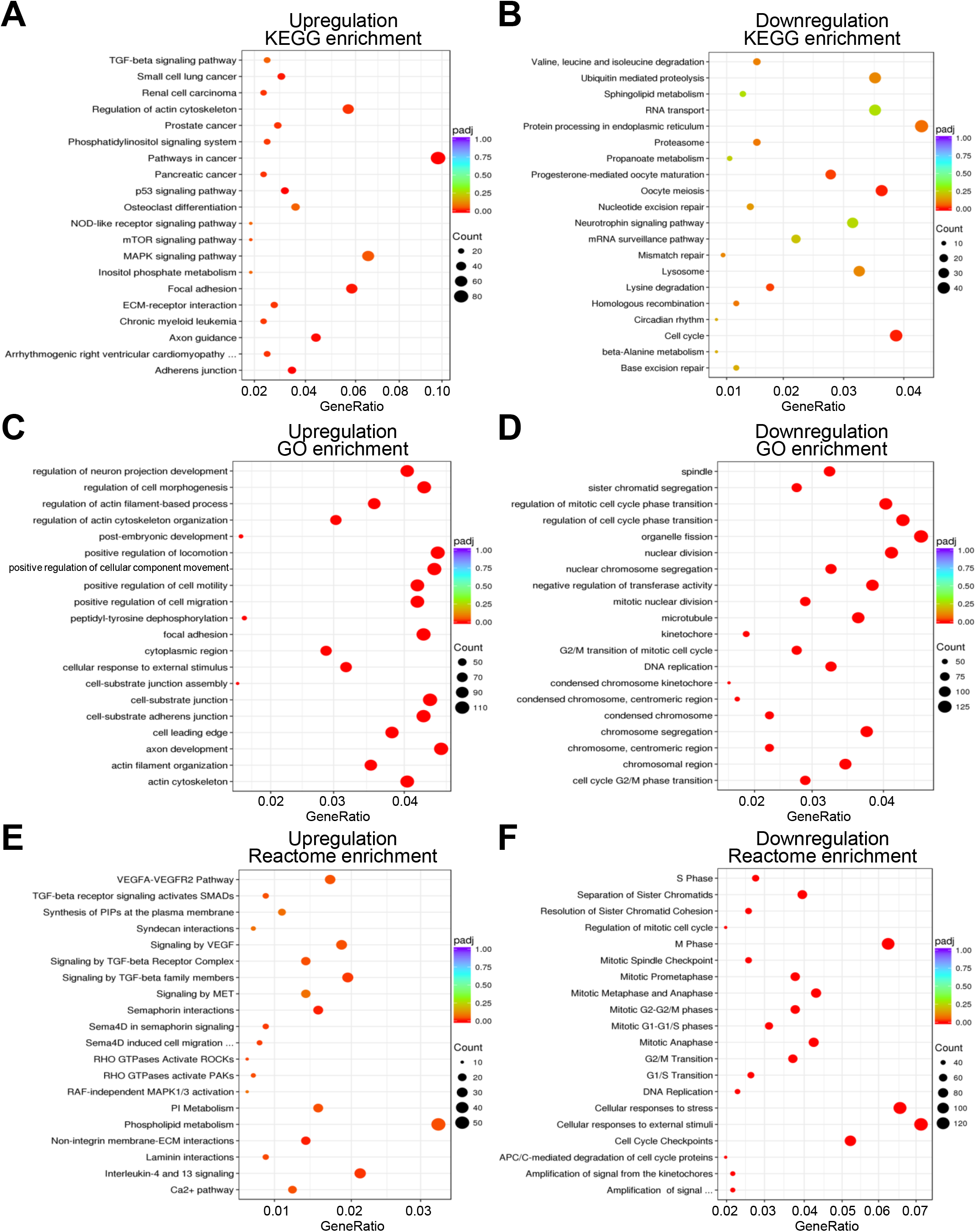
Functional analysis of the differential expressed genes by FoxQ1 overexpression in SUM159 cells. ***A-B*,** scatter plots for Kyoto Encyclopedia of Genes and Genomes (KEGG) pathway analyses showing upregulation (***A***) or downregulation (***B***) by FoxQ1 overexpression. ***C-D*,** scatter plots for gene ontology (GO) enrichment analyses showing upregulation (***C***) or downregulation (***D***) by FoxQ1 overexpression. ***E*-*F*,** scatter plots for Reactome pathway analyses showing upregulation (***E***) or downregulation (***F***) by FoxQ1 overexpression. padj, adjusted *p*-value.

### The gene ontology (GO) and Reactome pathway analyses

All three ontologies including cellular component, molecular function, and biological processes are included in the GO enrichment analysis. The GO enrichment analysis revealed that FoxQ1 overexpression caused upregulation of genes associated with positive regulation of locomotion, axon development, positive regulation of cellular component movement, cell substrate/adherens junction, focal adhesion, positive regulation of cell motility/migrations and a few other pathways (Fig. 2*C*). The downregulated genes following FoxQ1 overexpression in SUM159 cells were mostly associated with the GO pathway terms related to cell cycle regulation (GO terms: regulation of cell cycle phase transition, regulation of mitotic cell cycle phase transition, nuclear division, nuclear chromosome segregation, microtubule, cell cycle G2/M phase transition, centromeric region, G2/M transition of mitotic cell cycle, etc.) (Fig. *2D*).

The Reactome database contains annotations for diverse set of molecular and cell biological topics such as cell cycle, metabolism, signaling, transport, cell motility, immune function, host-virus interaction, and neural function. Genes associated with phospholipid metabolism, VEGF/VEGF receptor pathway, transforming growth factor β pathway were significantly upregulated in FoxQ1 overexpressing SUM159 cells (Fig. *2E*). Like KEGG and GO pathway analyses, most downregulated genes from the Reactome analyses were associated with cell cycle regulation (Fig. *2F*). Genes associated with cellular response to stress/external stimuli were also significantly downregulated by FoxQ1 overexpression (Fig. 2*F*).

### Comparison of RNA-Seq data with published literature

Published studies have revealed multiple downstream targets of FoxQ1 in breast cancer (7–12). For example, Zhang *et al*. (7) showed downregulation of CDH1 (E-cadherin) but upregulation of mesenchymal markers including CTNNA1 (catenin alpha 1), CDH2 (N-cadherin), and FN1 (fibronectin 1) in highly metastatic breast cancer cell lines (e.g., MDA-MB-435, MDA-MB-231, SUM149, etc.) in comparison with breast cancer cell lines with low metastatic capacity (e.g., MCF-7, MDA-MB-361, BT20, etc.). As shown in Fig. *3A*, the RNA-seq results from the present study were consistent with the findings of Zhang *et al*. (7). Meng *et al*. (9) showed downregulation of *CST6, SEMA3A, ADAM9, THBS1, FOXA1*, and *EDN1* but upregulation of *PDGFRA, JAM3*, and *ZEB2* in FoxQ1 overexpressing human mammary epithelial cell line (HMLE) when compared to EV cells, and these changes were also observed in the RNA-Seq data in the present study (Fig. 3*B*). We have shown previously that promotion of stem-like phenotype in FoxQ1 overexpressing SUM159 cell line is associated with suppression of *DACH1* and *ZEB1* expression and upregulation of mRNA levels of *MYC* and *TWIST2* (10) and RNA-Seq results from the present study were consistent with these published observations (Fig. 3*C*). At the same, some published changes in genes downstream of FoxQ1 (7–9,11) were not validated in the RNA-Seq data from the present study, including, *PLD1*, *SNAI1*, *CTNNB1*, *JUP*, *CD44, S100A4. DCN, PDGFRB, CADM3, COL1A1*, *COL6A1*, *DSG2*, and *TWIST1* (Fig. 4*A-D*). Two possibilities exist to explain the inconsistencies between published data and RNA-Seq results. The RNA-Seq results were obtained using SUM159 cells and this cell line was not used in other published studies (8,9) except for our own published study (10). Secondly, the possibility of regulatory functions of other transcription factors can’t be excluded.

**Figure 3.**
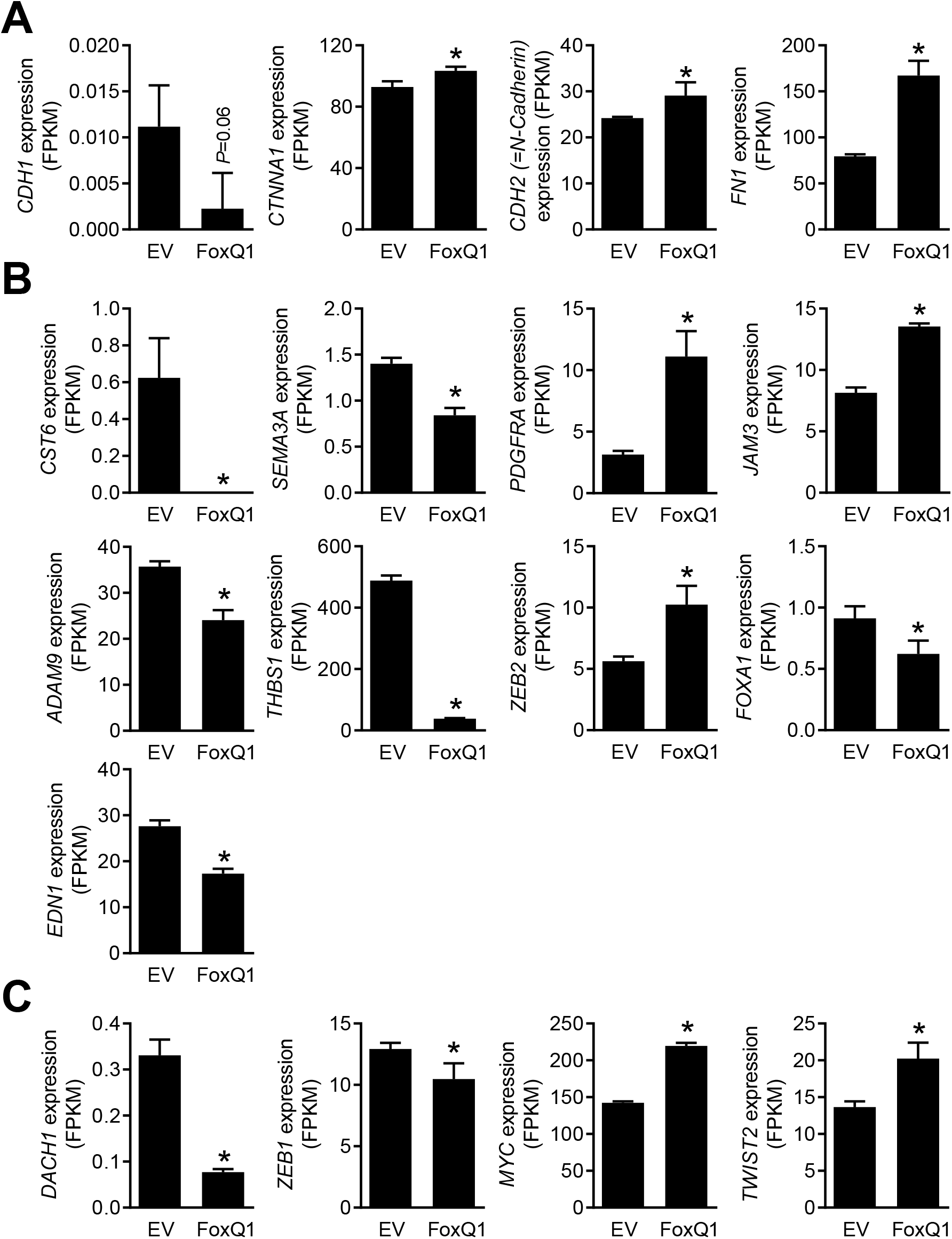
Consistent data from RNA-Seq analysis (present study) and published literature (7.9.10) on known targets of FoxQ1. ***A*,** RNA-seq analysis of genes associated with epithelial-mesenchymal transition (7). *CDH1*, *E-cadherin; CTNNA1, catenin alpha 1; CDH2, N-cadherin; FN1, fibronectin 1*. ***B*,** RNA-seq analysis of genes involved in breast cancer stemness and chemoresistance (9). *CST6*, *cystatin 6; SEMA3A*, *semaphorin 3A*; *PDGFRA*, *platelet derived growth factor receptor alpha*; *JAM3*, *junctional adhesion molecule 3*; *ADAM9*, *ADAM metallopeptidase domain 9*; *THBS1*, *thrombospondin 1*; *ZEB2*, *zinc finger E-box binding homeobox 2*; *FOXA1*, *forkhead box A1; EDN1*, *endothelin 1. **C***, RNA-seq analysis of genes related to breast cancer stemness (10). *DACH1, dachshund homolog 1; ZEB1, zinc finger E-box binding homeobox 1; TWIST2, twist basic helix-loop-helix transcription factor 2*. Results shown are mean ± S.D. (*n* = 3). *P < 0.05 by two-sided Student’s t-test.

**Figure 4.**
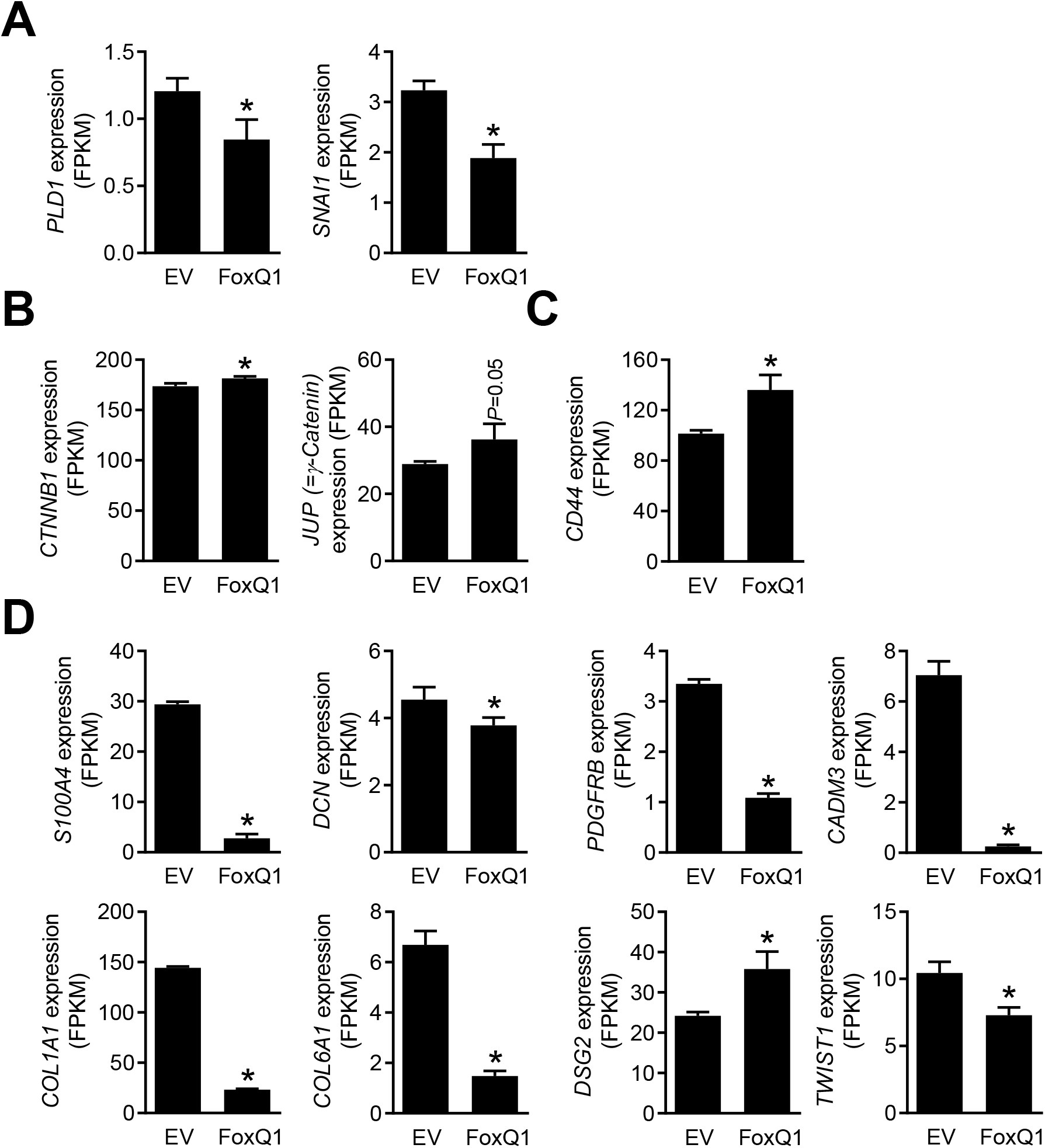
Inconsistent data between RNA-Seq analysis (present study) and published literature (7–9,11). ***A-D*,** RNA-seq analysis of genes not consistent with the published literatures (7,9,11). Results shown are mean ± S.D. (*n* = 3). *P < 0.05 by two-sided Student’s t-test. *PLD1, phospholipase D1; SNAI1, snail family transcriptional repressor 1; CTNNB1, catenin beta 1; S100A4, S100 calcium binding protein A4; DCN, Decorin; PDGFRB, platelet derived growth factor receptor beta; CADM3, cell adhesion molecule 3; COL1A1, collagen type I alpha 1; COL6A1, collagen Type VI alpha 1; DSG2, desmoglein 2; TWIST1, twist basic helix-loop-helix transcription factor 1*.

### Effect of FoxQ1 overexpression of cell cycle progression

Flow histograms for cell cycle distribution in EV and FoxQ1 overexpressing SUM159 cells after serum starvation and collection at different time points after release into the complete medium are shown in Fig. 5*A*. As expected, serum starvation caused accumulation of G0/G1 phase fraction in both cell lines. The FoxQ1 overexpressing SUM159 cells, but not MCF-7 cells, exhibited a significant decrease in fraction of G0/G1 phase cells that was accompanied by S and G2/M phase cell cycle arrest (Fig. 5*B*). These results indicated S and G2/M phase cell cycle arrest upon FoxQ1 overexpression in SUM159 cells but not in MCF-7 cells.

**Figure 5.**
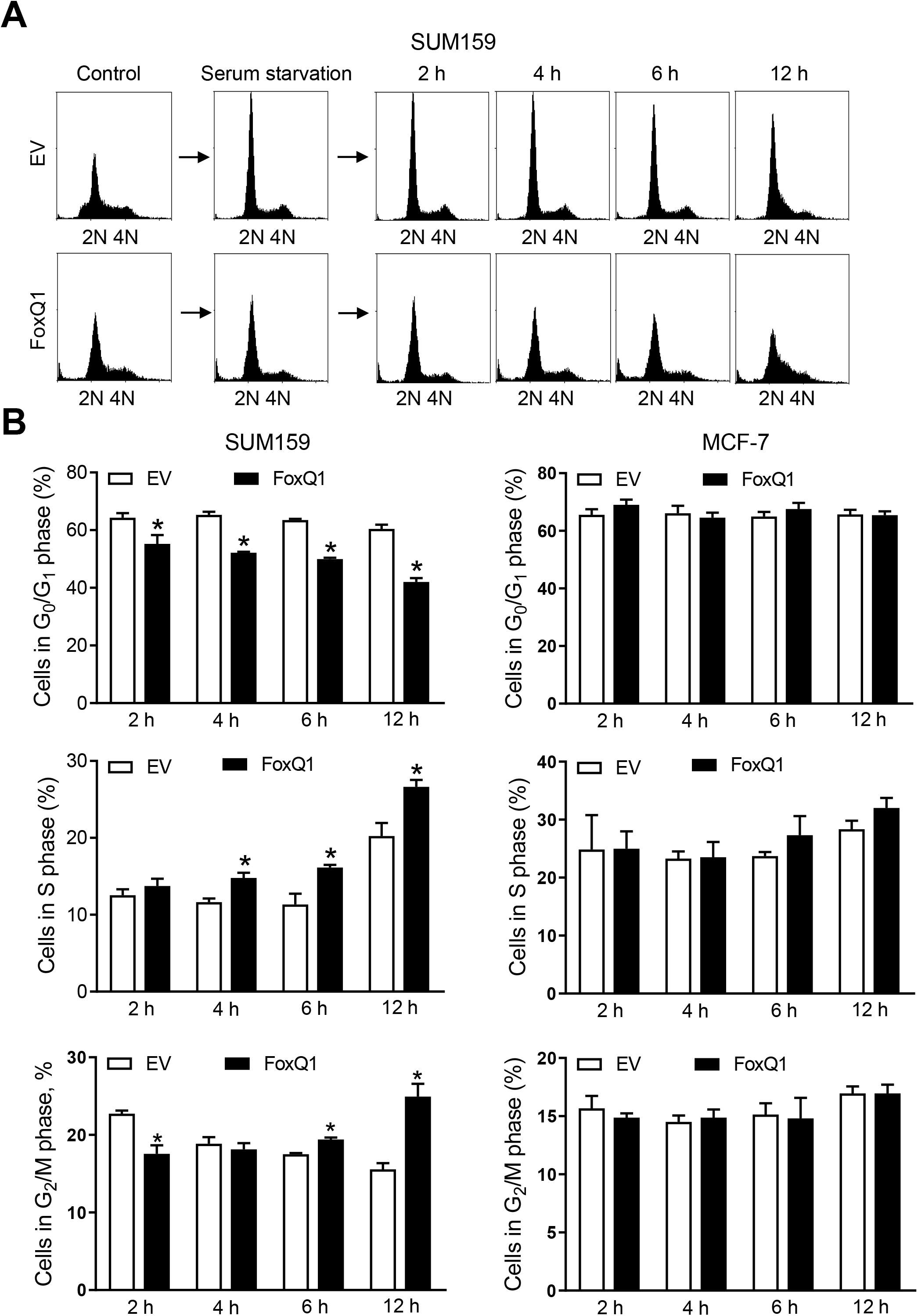
FoxQ1 overexpression resulted in cell cycle arrest at S and/or G2/M phases in SUM159 cells, but not in MCF-7 cells. ***A*,** representative flow histograms showing serum-starved synchronization at G_0_/G_1_ phase and resumption of cell cycle after release of cells in complete medium. ***B*,** percentage of G_0_/G_1_, S, and G2/M population. Results shown are mean ± SD (*n* = 3). *P < 0.05 by two-sided Student’s *t* test. Experiments were done at least three times with similar results.

Next, we performed western blotting and flow cytometry to quantitate mitotic fraction in FoxQ1 overexpressing cells and corresponding EV cells. Opposite effects were observed in basal-like SUM159 cells and luminal-type MCF-7 cells. FoxQ1 overexpression in SUM159 cells resulted in increased Ser10 phosphorylation of histone H3, which is a marker of mitotic cells (Fig. 6*A*). In the MCF-7 cell line, Ser10 phosphorylation of histone H3 was significantly lower in FoxQ1 overexpressing cells than in EV cells (Fig. 6*A*). These results were confirmed by flow cytometry (Fig. 6*B*,*C*). Because of the cell line-specific differences, we also performed similar experiments in another basal-like cell line (HMLE). The mitotic arrest by FoxQ1 overexpression was also observed in the HMLE cells similar to SUM159 (Fig. 6 *B,C*). These results indicated different role for FoxQ1 in basal-like *versus* luminal-type breast cancer cells.

**Figure 6.**
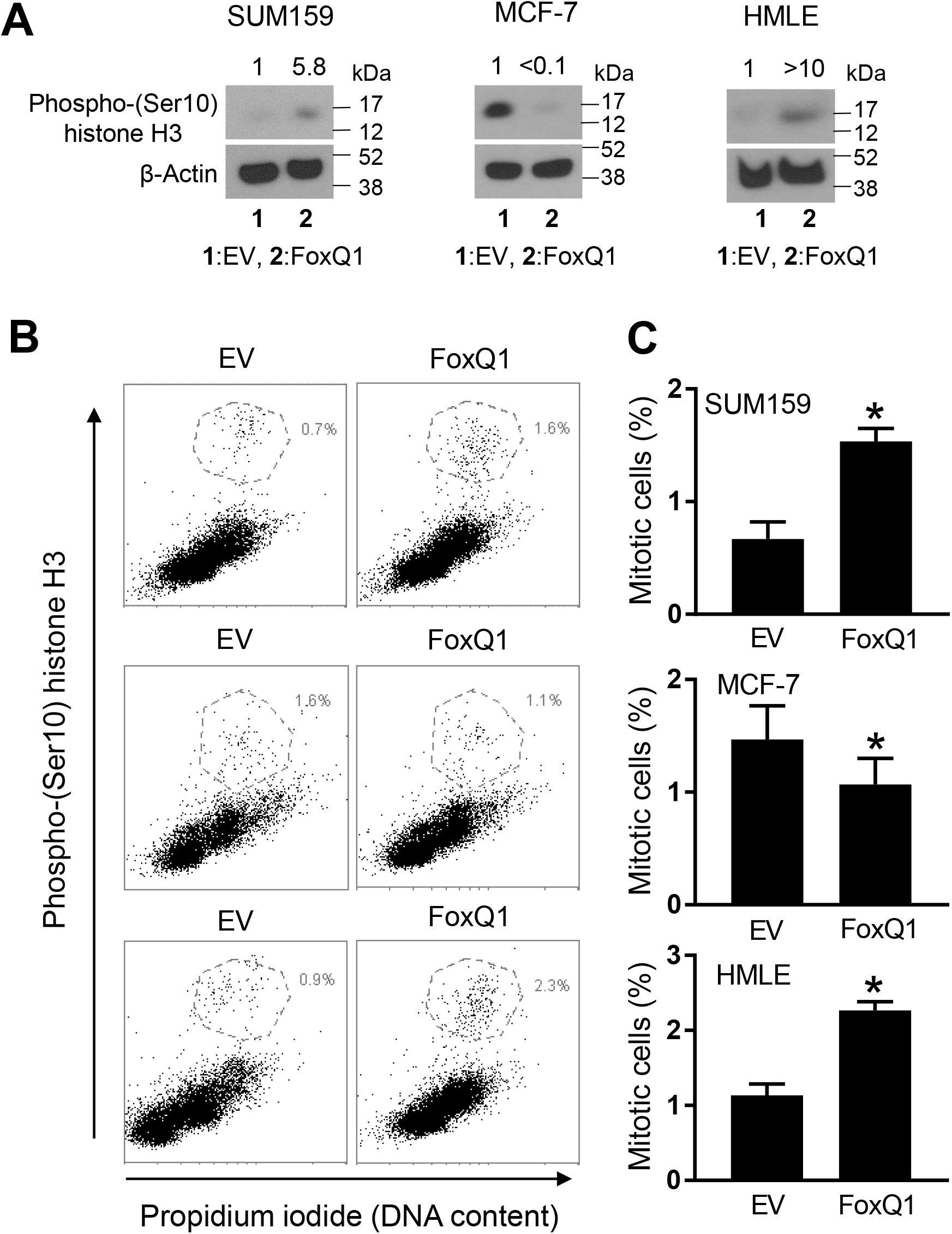
FoxQ1 overexpression resulted in mitotic arrest. ***A*,** Western blot analysis for phospho-(Ser10) histone H3 protein using lysates from (EV) cells or FoxQ1 overexpressing SUM159, MCF-7, and HMLE cells. The numbers on top of the bands represent changes in protein level compared to corresponding EV cells. ***B*,** representative flow histograms showing mitotic fraction in EV cells or FoxQ1 overexpressing SUM159, MCF-7, and HMLE cells. ***C*,** quantitation of mitotic fraction. Data shown are mean ± SD (*n* = 3-6). *P < 0.05 by two-sided Student’s t-test. Experiments were done twice with comparable results.

Overexpression of FoxQ1 in the MCF-7 cell line resulted in suppression of protein levels of cyclin-dependent kinase 2 (CDK2) and Cyclin D1, but upregulation of cell division cycle 25C (CDC25C), CDK4, and Cyclin B1 protein expression (Fig. 7). The protein levels of Cyclin A and CDK1 were not affected by FoxQ1 overexpression in the MCF-7 cell line (Fig. 7). The results were strikingly different in the SUM159 cell line stably transfected with FoxQ1 that exhibited downregulation of CDC25C, Cyclin B1, and CDK1 but upregulation of Cyclin D1, Cyclin A, and CDK4 (Fig. 7). Collectively, these results indicated cell line-specific responses upon FoxQ1 overexpression with respect to cell cycle progression.

**Figure 7.**
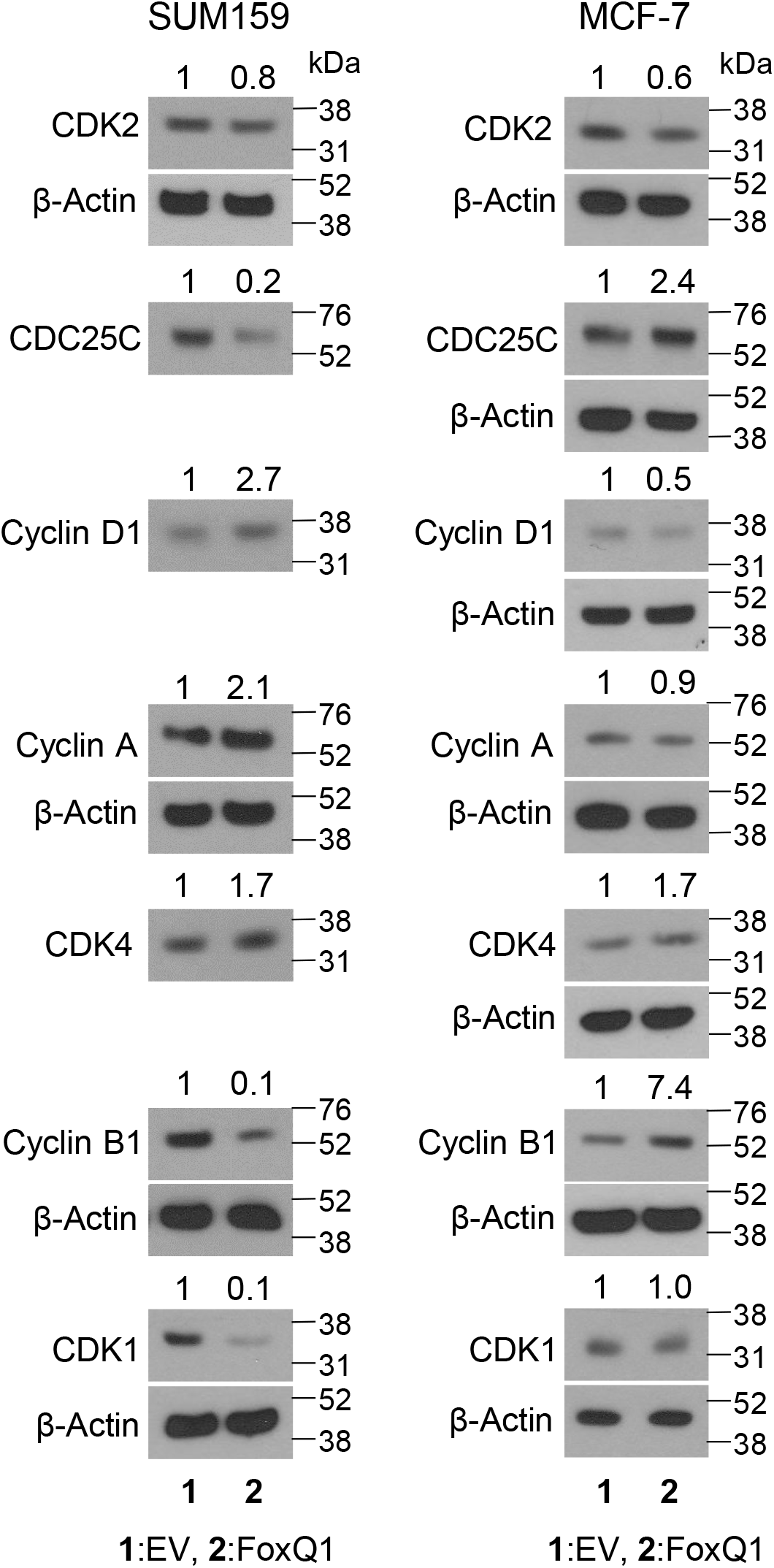
Overexpression of FoxQ1 modulated expression of proteins associated with cell cycle regulation. Representative immunoblots for CDK2, CDC25C, Cyclin D1, Cyclin A, CDK4, Cyclin B1, and CDK1 using lysate from EV or FoxQ1 overexpressing SUM159 or MCF-7 cells. Numbers above the bands are relative expression level of each protein compared to EV cells. The β-Actin blot is the same for some proteins. Experiment were done at least twice with comparable results. CDK2, cyclin dependent kinase 2; CDC25C, cell division cycle 25C; CDK4, cyclin dependent kinase 4; CDK1, cyclin dependent kinase 1.

### IL-1α, IL-8, and VEGF are novel downstream transcriptional targets of FoxQ1

The Reactome pathways analysis also showed downregulation of genes associated with cellular response to stress upon FoxQ1 overexpression in the SUM159 cell line (Fig. 2*F*). Table 2 lists genes associated with cellular response to stress whose expression was altered upon FoxQ1 overexpression. We compared the correlations observed from the RNA-Seq data (present study) with The Cancer Genome Atlas (TCGA) dataset. Positive correlation was observed for several upregulated or downregulated genes from the RNA-Seq data and the breast cancer TCGA analysis (Table 2). On the other hand, RNA-Seq data was not consistent with that of breast cancer TCGA for a few other genes (identified by negative correlation in Table 2). Because FoxQ1 has oncogenic function, we focused on IL-1α for further investigation. RNA-seq data indicated a 23-fold increase in expression of *IL-1α* in FoxQ1 overexpressing SUM159 cells compared with EV cells (Fig. 8*A*). The breast cancer TCGA analysis revealed a significant positive association between expression of *IL-1α* and that of *FoxQ1* (Fig. 8*B*). qRT-PCR confirmed overexpression of *IL-1α* mRNA upon forced expression of FoxQ1 in SUM159 cells (Fig. 8*C*). Secretion of IL-1α in the media was also significantly higher in the FoxQ1 overexpressing SUM159 cells when compared to EV cells (Fig. 8*D*). The promoter of *IL-1α* contains a single FoxQ1 binding consensus sequence of TGTTTA (9). Chromatin immunoprecipitation (ChIP) confirmed recruitment of FoxQ1 at the promoter of *IL-1α* in SUM159 cells (Fig. 8*E*). The expression of IL-1α was very low in MCF-7 cells when compared to SUM159 and thus these experiments were not performed in the former cell line. These results indicated that *IL-1α* is a direct transcriptional target of FoxQ1 at least in the SUM159 cell line.

**Table 2.**
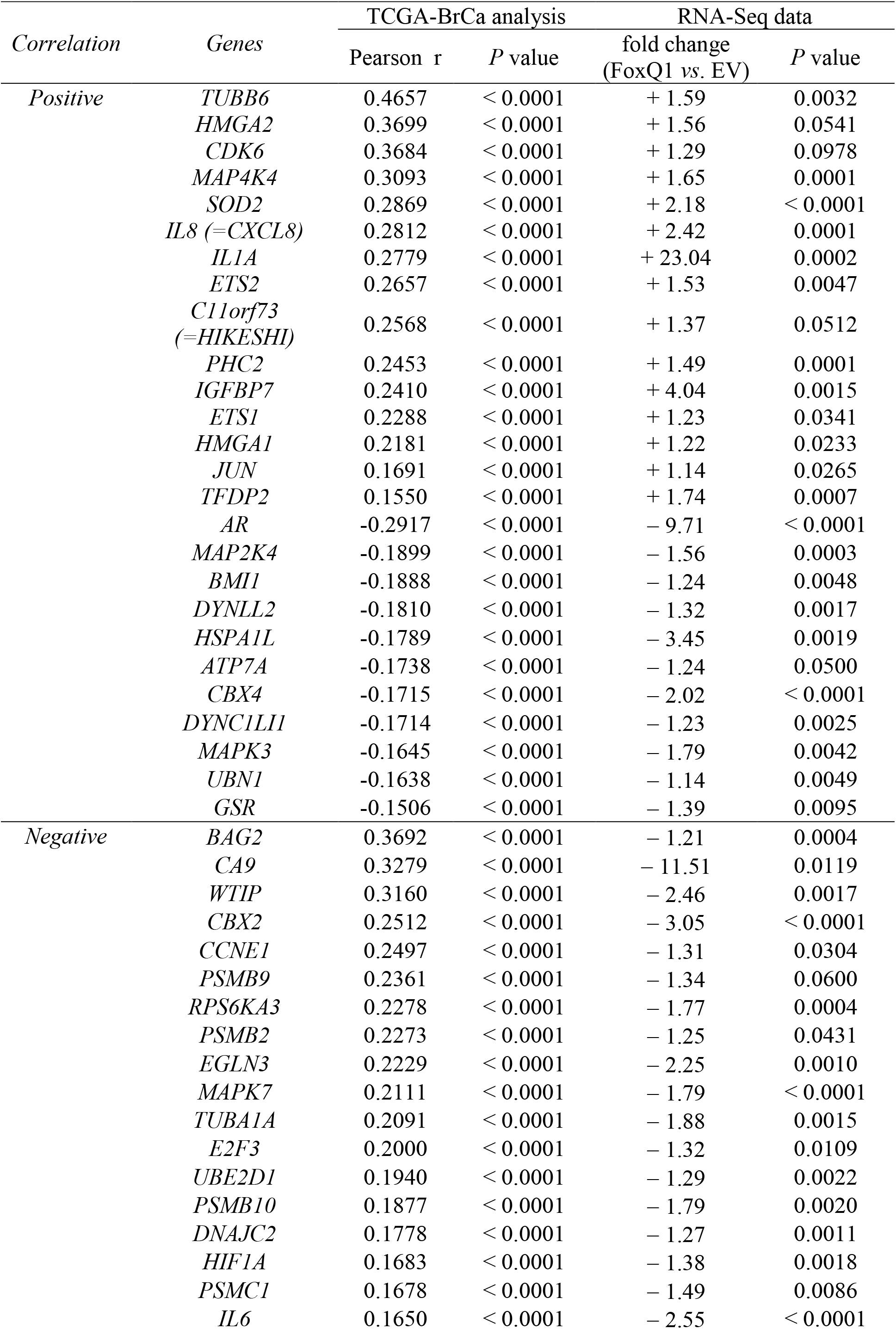
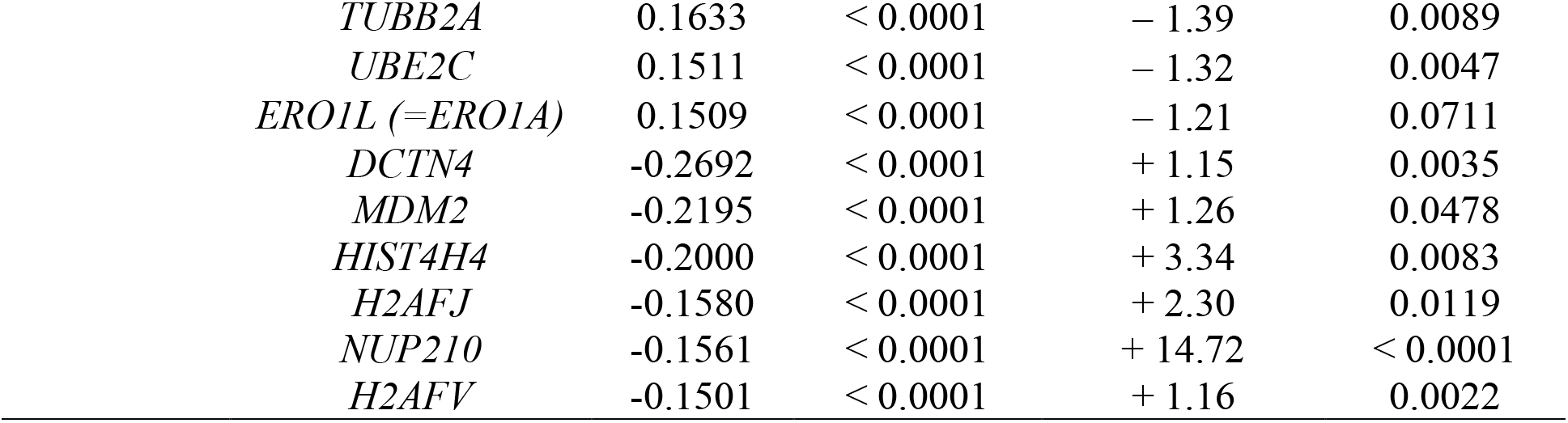
Correlation between *FoxQ1* and genes associated with “Cellular response to stress” category from TCGA-BrCa (n=1097) and RNA-seq results. Student *t* test was used for statistical analysis.

| *Correlation* | *Genes* | TCGA-BrCa analysis |  | RNA-Seq data |  |
| --- | --- | --- | --- | --- | --- |
|  |  | Pearson *r* | *P* value | fold change<br>(FoxQ1 vs. EV) | *P* value |
| *Positive* | *TUBB6* | 0.4657 | < 0.0001 | + 1.59 | 0.0032 |
|  | *HMGA2* | 0.3699 | < 0.0001 | + 1.56 | 0.0541 |
|  | *CDK6* | 0.3684 | < 0.0001 | + 1.29 | 0.0978 |
|  | *MAP4K4* | 0.3093 | < 0.0001 | + 1.65 | 0.0001 |
|  | *SOD2* | 0.2869 | < 0.0001 | + 2.18 | < 0.0001 |
|  | *IL8 (=CXCL8)* | 0.2812 | < 0.0001 | + 2.42 | 0.0001 |
|  | *IL1A* | 0.2779 | < 0.0001 | + 23.04 | 0.0002 |
|  | *ETS2* | 0.2657 | < 0.0001 | + 1.53 | 0.0047 |
|  | *C11orf73*<br>(=*HIKESHI*) | 0.2568 | < 0.0001 | + 1.37 | 0.0512 |
|  | *PHC2* | 0.2453 | < 0.0001 | + 1.49 | 0.0001 |
|  | *IGFBP7* | 0.2410 | < 0.0001 | + 4.04 | 0.0015 |
|  | *ETS1* | 0.2288 | < 0.0001 | + 1.23 | 0.0341 |
|  | *HMGA1* | 0.2181 | < 0.0001 | + 1.22 | 0.0233 |
|  | *JUN* | 0.1691 | < 0.0001 | + 1.14 | 0.0265 |
|  | *TFDP2* | 0.1550 | < 0.0001 | + 1.74 | 0.0007 |
|  | *AR* | -0.2917 | < 0.0001 | - 9.71 | < 0.0001 |
|  | *MAP2K4* | -0.1899 | < 0.0001 | - 1.56 | 0.0003 |
|  | *BMI1* | -0.1888 | < 0.0001 | - 1.24 | 0.0048 |
|  | *DYNLL2* | -0.1810 | < 0.0001 | - 1.32 | 0.0017 |
|  | *HSPA1L* | -0.1789 | < 0.0001 | - 3.45 | 0.0019 |
|  | *ATP7A* | -0.1738 | < 0.0001 | - 1.24 | 0.0500 |
|  | *CBX4* | -0.1715 | < 0.0001 | - 2.02 | < 0.0001 |
|  | *DYNC1LI1* | -0.1714 | < 0.0001 | - 1.23 | 0.0025 |
|  | *MAPK3* | -0.1645 | < 0.0001 | - 1.79 | 0.0042 |
|  | *UBN1* | -0.1638 | < 0.0001 | - 1.14 | 0.0049 |
|  | *GSR* | -0.1506 | < 0.0001 | - 1.39 | 0.0095 |
| *Negative* | *BAG2* | 0.3692 | < 0.0001 | - 1.21 | 0.0004 |
|  | *CA9* | 0.3279 | < 0.0001 | - 11.51 | 0.0119 |
|  | *WTIP* | 0.3160 | < 0.0001 | - 2.46 | 0.0017 |
|  | *CBX2* | 0.2512 | < 0.0001 | - 3.05 | < 0.0001 |
|  | *CCNE1* | 0.2497 | < 0.0001 | - 1.31 | 0.0304 |
|  | *PSMB9* | 0.2361 | < 0.0001 | - 1.34 | 0.0600 |
|  | *RPS6KA3* | 0.2278 | < 0.0001 | - 1.77 | 0.0004 |
|  | *PSMB2* | 0.2273 | < 0.0001 | - 1.25 | 0.0431 |
|  | *EGLN3* | 0.2229 | < 0.0001 | - 2.25 | 0.0010 |
|  | *MAPK7* | 0.2111 | < 0.0001 | - 1.79 | < 0.0001 |
|  | *TUBA1A* | 0.2091 | < 0.0001 | - 1.88 | 0.0015 |
|  | *E2F3* | 0.2000 | < 0.0001 | - 1.32 | 0.0109 |
|  | *UBE2D1* | 0.1940 | < 0.0001 | - 1.29 | 0.0022 |
|  | *PSMB10* | 0.1877 | < 0.0001 | - 1.79 | 0.0020 |
|  | *DNAJC2* | 0.1778 | < 0.0001 | - 1.27 | 0.0011 |
|  | *HIF1A* | 0.1683 | < 0.0001 | - 1.38 | 0.0018 |
|  | *PSMC1* | 0.1678 | < 0.0001 | - 1.49 | 0.0086 |
|  | *IL6* | 0.1650 | < 0.0001 | - 2.55 | < 0.0001 |

|  |  |  |  |  |
| --- | --- | --- | --- | --- |
| *TUBB2A* | 0.1633 | < 0.0001 | − 1.39 | 0.0089 |
| *UBE2C* | 0.1511 | < 0.0001 | − 1.32 | 0.0047 |
| *ERO1L (=ERO1A)* | 0.1509 | < 0.0001 | − 1.21 | 0.0711 |
| *DCTN4* | -0.2692 | < 0.0001 | + 1.15 | 0.0035 |
| *MDM2* | -0.2195 | < 0.0001 | + 1.26 | 0.0478 |
| *HIST4H4* | -0.2000 | < 0.0001 | + 3.34 | 0.0083 |
| *H2AFJ* | -0.1580 | < 0.0001 | + 2.30 | 0.0119 |
| *NUP210* | -0.1561 | < 0.0001 | + 14.72 | < 0.0001 |
| *H2AFV* | -0.1501 | < 0.0001 | + 1.16 | 0.0022 |

**Figure 8.**
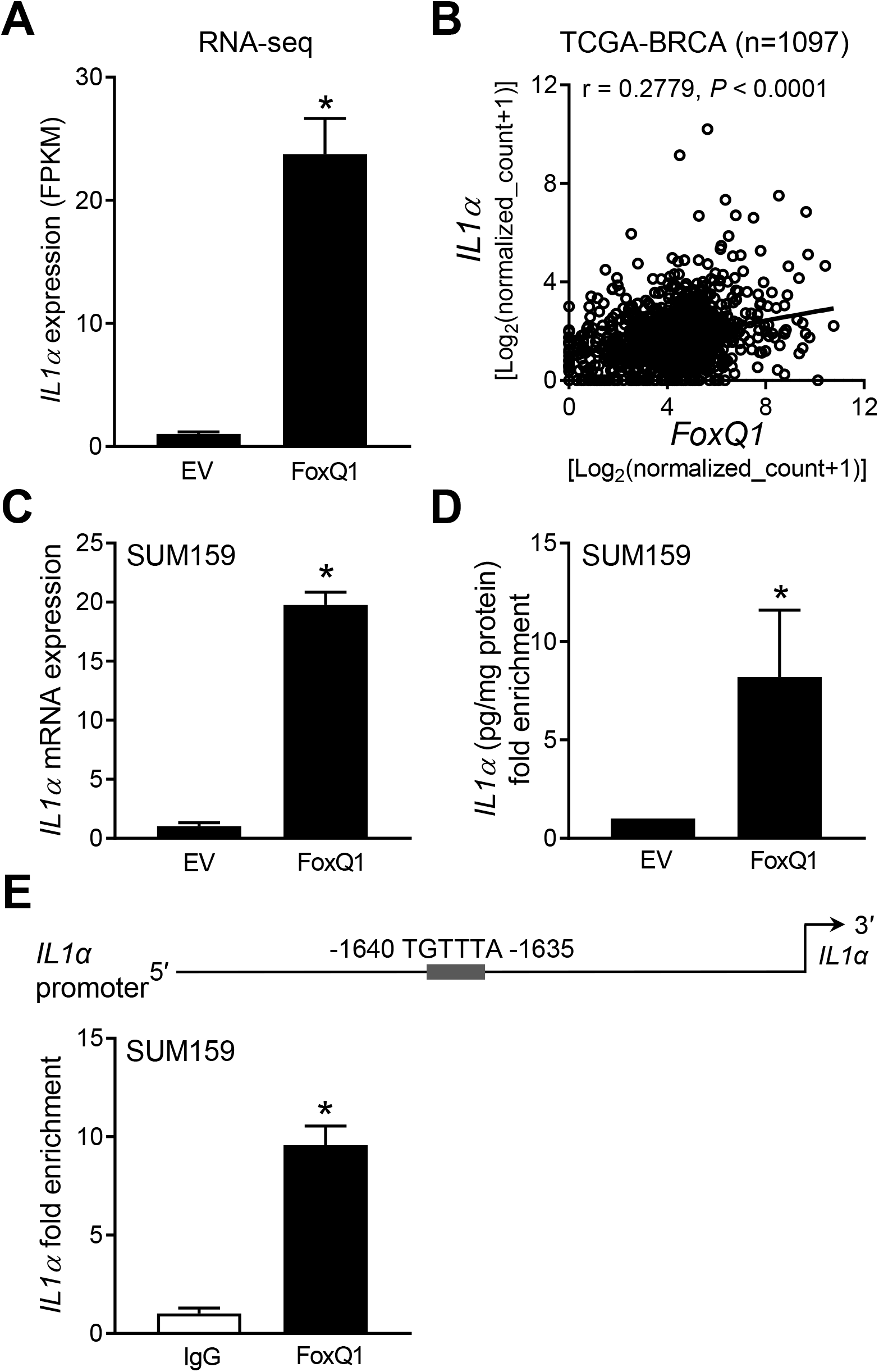
FoxQ1 is a novel regulator of *IL-1α*. ***A*,** RNA-seq analysis of *IL-1α* gene expression in EV cells FoxQ1 overexpressing SUM159 cells. Data shown are mean ± S.D. (*n* = 3). *P < 0.05 by two-sided Student’s t-test. ***B*,** positive correlation between *FoxQ1* and *IL-1α* expression in breast tumor TCGA dataset (*n* = 1097). Pearson’s test was used to determine statistical significance of the correlation. ***C*,** quantification of *IL-1α* mRNA expression in EV and FoxQ1 overexpressing SUM159 cells. Data shown are mean ± S.D. (*n* = 3). *P < 0.05 by two-sided Student’s t-test. ***D*,** quantification of IL-1α secretion in the media of EV or FoxQ1 overexpressing SUM159 cells. Data shown are mean ± S.D. (*n* = 3). *P < 0.05 by two-sided Student’s t-test. ***E*,** identification of a putative FoxQ1 binding site in the promoter region of *IL-1α*. The bar graph shows recruitment of FoxQ1 at the *IL-1α* promoter by ChIP analysis. The results shown are mean ± S.D. (*n* = 3). *P < 0.05 by two-sided Student’s t-test. Experiments were done twice with similar results.

The RNA-seq data indicated a significantly higher level of *IL-8* mRNA in FoxQ1 overexpressing SUM159 cells compared to EV cells (Fig. 9*A*). A significant positive association between expression of *IL-8* and that of *FoxQ1* was also observed in the breast cancer TCGA dataset (Fig. 9*B*). Overexpression of *IL-8* in FoxQ1 overexpressing SUM159 cell was confirmed by qRT-PCR (Fig. 9*C*). The level of IL-8 in the media was also significantly higher in the FoxQ1 overexpressing SUM159 cells when compared to EV cells (Fig. 9*D*). Three FoxQ1 binding consensus sequences were observed at the promoter of *IL-8*. ChIP assay confirmed recruitment of FoxQ1 at two of those sites of *IL-8* promoter (Fig. 9*E*). Similarly, FoxQ1 overexpressing MCF-7 cells exhibited a significantly higher level of IL-8 secretion in comparison with EV cells (Fig. 10*A*). Furthermore, FoxQ1 was recruited at two sites of the *IL-8* promoter in MCF-7 cells (Fig. 10*B*).

**Figure 9.**
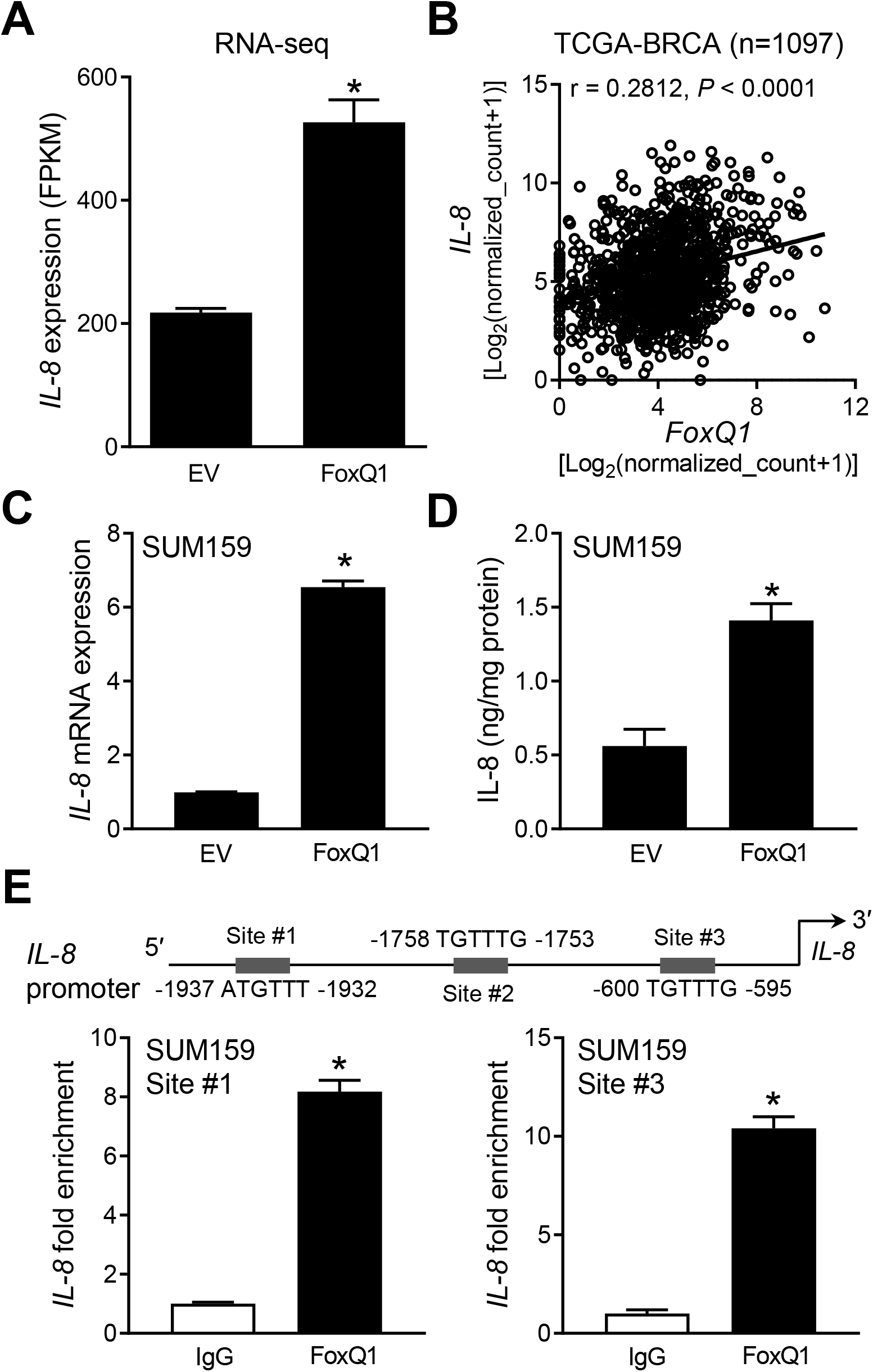
IL-8 is a novel downstream target of FoxQ1 in SUM159 cells. ***A*,** RNA-seq analysis for *IL-8* gene expression in EV cells and FoxQ1 overexpressing SUM159 cells. Data shown are mean ± S.D. (*n* = 3). *P < 0.05 by two-sided Student’s t-test. ***B*,** positive correlation between *FoxQ1* and *IL-8* expression in breast tumor TCGA dataset (*n* = 1097). Pearson’s test was used to determine statistical significance of the correlation. ***C*,** quantification of *IL-8* mRNA expression by qRT-PCR in EV and FoxQ1 overexpressing SUM159 cells. Data shown are mean ± S.D. (*n* = 3). *P < 0.05 by two-sided Student’s t-test. ***D*,** quantification of IL-8 secretion in the media of EV and FoxQ1 overexpressing SUM159 cells. Data shown are mean ± S.D. (*n* = 3). *P < 0.05 by two-sided Student’s t-test. ***E*,** identification of putative FoxQ1 binding sites at the promoter region of *IL-8*. The bar graph shows recruitment of FoxQ1 at the *IL-8* promoter by ChIP analysis. The results shown are mean ± S.D. (*n* = 3). *P < 0.05 by two-sided Student’s t-test. Experiments were repeated with similar results.

**Figure 10.**
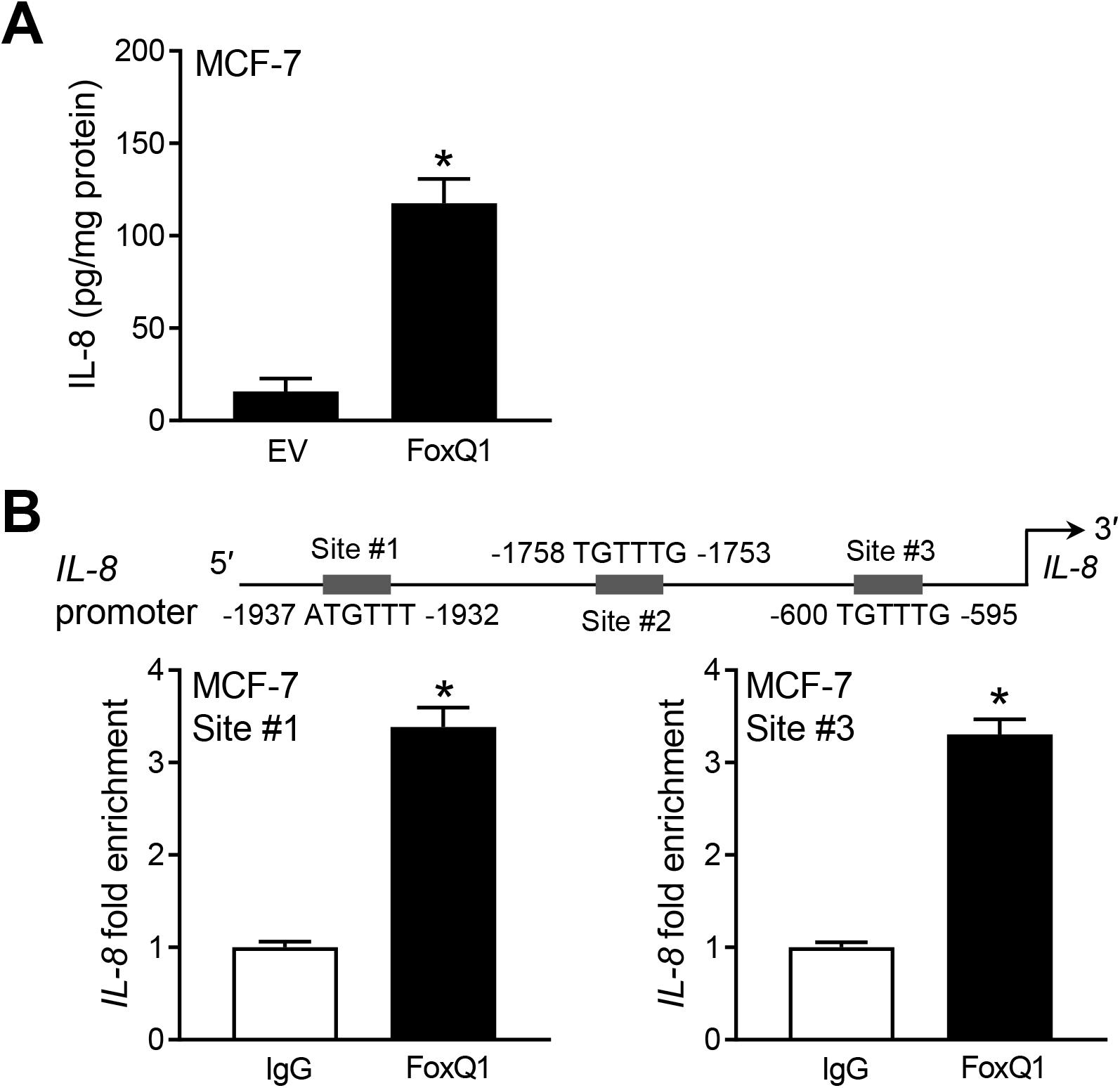
FoxQ1 regulated *IL-8* expression in MCF-7 cells. ***A*,** quantification of IL-8 secretion in the media of EV and FoxQ1 overexpressing MCF-7 cells. Data shown are mean ± S.D. (*n* = 3). *P < 0.05 by two-sided Student’s *t* test. ***B*,** ChIP assay for FoxQ1 recruitment at the promoter region of *IL-8*. The results shown are mean ± S.D. (*n* = 3). *P < 0.05 by two-sided Student’s t-test. Experiments were done twice with similar results.

The Reactome pathway analysis also revealed upregulation of genes associated with VEGF/VEGFR2 pathway in FoxQ1 overexpressing SUM159 cells. The RNA-Seq data revealed upregulation of *VEGFA* mRNA in FoxQ1 overexpressing SUM159 cells when compared to EV cells (Fig. 11*A*). Analysis of the breast cancer TCGA dataset also revealed a positive correlation between expression of *FoxQ1* and that of *VEGF* (Fig. 11*B*). The *VEGF* mRNA level (Fig. 11*C*) and/or secretion of the protein (Fig. 11*C*) were increased upon FoxQ1 overexpression. ChIP assay indicated recruitment of FoxQ1 at all 3 sites of *VEGFA* (Fig. 11*E*). Because of weak association between expression of FoxQ1 and VEGFA in MCF-7 cells, ChIP assay was not performed in this cell line.

**Figure 11.**
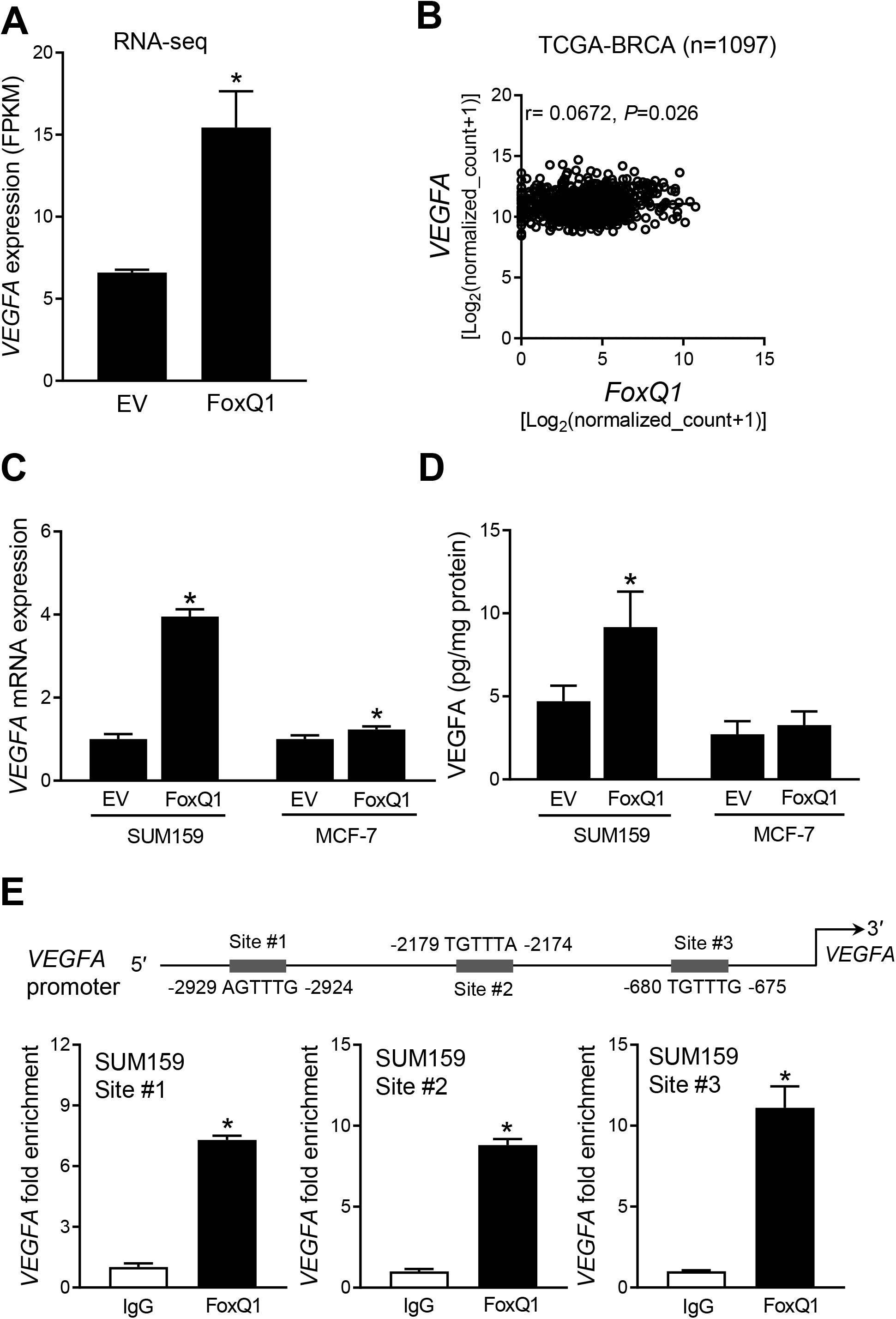
FoxQ1 regulates VEGFA expression in SUM159 cells. ***A*,** RNA-seq analysis of *VEGFA* gene expression difference between EV and FoxQ1 overexpressing SUM159 cells. Data shown are mean ± S.D. (*n* = 3). *P < 0.05 by two-sided Student’s *t* test. ***B*,** TCGA dataset showed the correlation between *FoxQ1* and *VEGFA* in breast tumor (n = 1097). Pearson’s correlation coefficient was used for the analysis. ***C*,** quantification of *VEGFA* mRNA expression by quantitative real-time PCR in EV or FoxQ1 overexpressing SUM159 and MCF-7 cells. Data shown are mean ± S.D. (*n* = 3). *P < 0.05 by two-sided Student’s *t* test. Experiments were done twice with comparable results. ***D*,** quantification of VEGFA secretion in EV or FoxQ1 overexpressing SUM159 and MCF-7 cells after 16 hours of serum-starvation. Data shown are mean ± S.D. (*n* = 3). *P < 0.05 by two-sided Student’s *t* test. Experiments were done twice with comparable results. *E*, identification of putative FoxQ1 binding sites in the promoter region of *VEGFA*. The bar graph shows recruitment of FoxQ1 in the *VEGFA* promoter by ChIP analysis. The results shown are mean ± S.D. (*n* = 3). *P < 0.05 by two-sided Student’s *t* test. Experiments were done twice with similar results.

## Discussion

FoxQ1, also known as HNF-3/fkh homolog-1 or HFH-1, belongs to the forkhead family of transcription factors and its normal physiological functions include regulation of hair differentiation and control of mucin expression and granule content in stomach surface mucous cells (12–15). The *FoxQ1* gene is located at chromosome 6p25.3 that encodes a 403-amino acid protein (16,17). Structurally, the FoxQ1 is characterized by alanine-and glycine-rich region, the forkhead box domain (Winged-helix or DNA-binding domain), and proline-rich region (17). The FoxQ1 is an evolutionary conserved protein with 100% amino acid sequence similarity in the DNA binding domain between human, mouse, and rat (17,18). Overexpression of protein or mRNA levels of FoxQ1 has been reported in different cancers, including gastric cancer, hepatocellular carcinoma, laryngeal carcinoma, pancreatic cancer, colorectal cancer, and breast cancer (8, 10, 19–24). We showed previously that the level of FoxQ1 protein is higher by 18.4-fold in basal-like human breast cancers compared to normal mammary tissue (10). Because luminal-type disease accounts for majority of human breast cancers, in this study we analyzed the level of FoxQ1 protein in this subtype. Like basal-like breast cancers, the expression of FoxQ1 protein was significantly higher in the luminal-type human breast cancers in comparison with normal breast tissues. These results suggest that oncogenic function of FoxQ1 may span across disease subtypes in human breast cancers.

In breast cancer, FoxQ1 is well-known for its role in promotion of EMT, cell migration and invasion, self-renewal of breast cancer stem like cells, and metastasis (7–10). Consistent with these published results (7–10), the GO pathways analysis of the RNA-Seq data also indicated upregulation of genes associated with positive regulation of locomotion, positive regulator of cellular component movement, positive regulation of cell motility, and positive regulation of cell migration. At the same time, the RNA-Seq data also suggests additional functions of FoxQ1 based on GO and Reactome pathway analyses, including regulation of cell morphogenesis, regulation of actin cytoskeleton organization, peptidyl-tyrosine dephosphorylation, and so forth. Further work is necessary to systematically test the role of FoxQ1 in these processes.

Studies have revealed direct regulation of multiple cancer-relevant genes by FoxQ1 (7–10). FoxQ1 functions to repress E-cadherin expression in breast cancer cell lines (8). For example, the promoter activity of E-cadherin was repressed by overexpression of FoxQ1 in HMLER cells but restored by knockdown of FoxQ1 in MDA-MB-231 cells (8). Another study confirmed these findings in 293FT and MCF-7 cells (7). The three regulatory E-boxes of the *E-cadherin* promoter were shown to cooperate in its repression mediated by FoxQ1 overexpression (7). Additional direct targets of FoxQ1 in breast cancer include transcription factors (Twist1 and Zeb2) and PDGFRα/β growth factor receptors to confer resistance to chemotherapy drugs like doxorubicin, paclitaxel, and imatinib *in vitro* and *in vivo* (9). The DACH1 homolog gene was originally identified as a critical regulator of eye and limb development in *Drosophila* (25). Studies have pointed to a tumor suppressor function of DACH1 in breast cancer (26–30). For example, overexpression of DACH1 inhibited breast cancer stem-like fraction *in vitro* and *in vivo* (27). Colony formation *in vitro* and xenograft growth *in vivo* was also significantly suppressed by DACH1 overexpression in MDA-MB-231 cells (26). DACH1 was also shown to inhibit EMT in breast cancer cells *in vitro* and repress metastatic potential of 4T1/Luc murine breast carcinoma cells *in vivo* in syngeneic mice (28). Our own group was first to implicate FoxQ1 in direct transcriptional repression of *DACH1* (10). In one study, DACH1 was shown to repress IL-8 level (30). Collectively, these studies indicate that FoxQ1 is a druggable target in breast and other cancers but a selective inhibitor of this transcription factor is yet to be identified.

We found that downregulation of genes associated with cell cycle regulation was the most prominent pathway enrichment by FoxQ1 overexpression from KEGG, GO, and Reactome pathway analyses (Fig. 2). We also found alterations in expression of many key proteins of the cell cycle regulation by FoxQ1 overexpression, but there are differences between SUM159 and MCF-7 cells. Consequently, FoxQ1 overexpression in SUM159 cells, but not in MCF-7, results in S, G2M and mitotic arrest. From the cell cycle distribution changes at least in SUM159 cells, one would expect a pronounced effect of FoxQ1 on cell proliferation and cell survival but the published *in vitro* data do not support this possibility at least in breast cancer (7,8). To the contrary, overexpression of FoxQ1 in non-small cell lung cancer cell lines (A549 and HCC827) results in increased proliferation when compared to corresponding EV cells (31). Cell proliferation is decreased upon FoxQ1 knockdown in a laryngeal carcinoma cell line (21). Because the full spectrum of the FoxQ1-regulated transcriptome is still not fully appreciated, further work may shed light as to why FoxQ1 overexpression is unable to increase breast cancer cell proliferation.

The present study, for the first time demonstrates that *IL-1α, IL-8*, and *VEGFA* are direct transcriptional targets of FoxQ1. FoxQ1 is recruited at the promoters of *IL-1α*, *IL-8*, and *VEGFA* genes. Moreover, protein levels of IL-1α, IL-8, and/or VEGFA are increased by FoxQ1 overexpression in SUM159 and/or MCF-7 cells. In breast cancer patients, IL-1α protein secretion is correlated with malignant phenotype (32). IL-1α-derived from cancer cells was shown to promote autocrine and paracrine induction of pro-metastatic genes in breast cancer cells (33). Interestingly, IL-1α protein expression in breast cancer correlated with that of IL-8 (34), and both these cytokines are direct targets of FoxQ1 (present study). Similar to the IL-1α, studies have established a pro-tumorigenic function of IL-8 in breast cancer (35–40). For example, the IL-8 expression in breast cancer cells is proportional to their invasion potential (37,38). The invasiveness of breast cancer cells is increased by overexpression of IL-8 as well as treatment with recombinant IL-8 (37,38). RNA interference of IL-8 in MDA-MB-231 cells diminishes its ability to invade (39). Studies have also indicated that IL-8 can promote stemness and EMT in breast cancer cells (41). These results indicate that promotion of EMT and stem-like phenotypes in breast and, possibly other cancers by FoxQ1 overexpression is partly mediated by direct regulation of the IL-1α and/or IL-8 expression.

More than 40,000 American women succumb to metastatic breast cancer every year (1). Increased tumor angiogenesis is considered critical for tumor growth (42,43). The VEGFA, routinely known as VEGF, is one of the major mediators of tumor neo-angiogenesis (44). The VEGF binds to VEGF receptor to promote vascular endothelial growth, migration, survival, and lymphangiogenesis (44,45). The present study shows that VEGFA is a direct transcriptional target of FoxQ1. However, this regulation seems more pronounced in the SUM159 than in MCF-7 cells. Thus, it is reasonable to conclude that VEGFA regulation by FoxQ1 is partly responsible for its pro-migratory and metastatic effects.

In conclusion, the present study reveals novel targets and functions of FoxQ1 in breast cancer. The RNA-Seq data also reveals additional direct regulatory targets of FoxQ1 at least in SUM159 cells that require further verification. For example, the Reactome pathways analysis of the RNA-Seq data from SUM159 cells indicates upregulation VEGF/VEGFR2 pathway, which plays an important role in endothelial cell proliferation and capillary tube formation (42), suggesting that this transcription factor can promote tumor angiogenesis.

## Experimental Procedures

### Reagents and cell lines

Cell culture reagents including fetal bovine serum, cell culture media, phosphate-buffered saline (PBS), and antibiotic mixture were purchased from Life Technologies-Thermo Fisher Scientific (Waltham, MA). Antibodies against CDK2, CDK4, Cyclin A, Cyclin D1, CDC25C, Cyclin B1, CDK1, and FoxQ1 (for chromatin immunoprecipitation assay) were from Santa Cruz Biotechnology (Dallas, TX). An antibody specific for detection of phospho- (Ser10) histone H3 was from Cell Signaling Technology (Danvers, MA). Anti-β-Actin antibody was from Sigma-Aldrich (St. Louis, MO).

The MCF-7 cell line was purchased from the American Type Culture Collection and authenticated by us in 2015 and 2017. The SUM159 cell line was purchased from Asterand (Detroit, MI) and authenticated by us in 2015 and 2017. Details of stable transfection of MCF-7 and SUM159 cells with pCMV6 empty vector and the same vector encoding FoxQ1 and their culture conditions have been described by us previously (10). The HMLE cells stably transfected with FoxQ1 and EV cells were generously provided by Dr. Guojun Wu (Karmanos Cancer Institute, Departments of Oncology and Pathology, Wayne State University, Detroit, MI) and maintained as recommended by the provider (7).

### Immunohistochemistry for FoxQ1 protein expression in tissue microarrays

Expression of FoxQ1 protein in tissue microarrays of human luminal-type breast cancers (US Biomax, Rockville, MD; catalog # BR1508) and normal human mammary tissues (US Biomax, Rockville, MD; catalog # BRN801a) was determined by immunohistochemistry essentially as described by us previously (10). At least three randomly-selected and non-overlapping fields on each core of the tissue microarray were examined using nuclear algorithm of the Aperio ImageScope software (Leica Biosystems, Buffalo Grove, IL). Data is expressed as H-score that is based on intensity (0, 1+, 2+, and 3+) % positivity (0-100%) and calculated using the following formula: (% of negative cells × 0) + (% of 1+ cells ×1) + (% of 2+ cells ×2) + (% of 3+ cells ×3). Some specimens for normal breast tissue (n=10) and luminal-type breast cancer (n=2) were not available for analysis due to poor staining and/or less than optimal sectioning on the tissue microarray (cracked or folded).

### RNA-seq analysis

FoxQ1 overexpressing SUM159 cells and EV cells (n=3 for each) were used for RNA-Seq analysis. Prior to RNA-Seq analysis, overexpression of FoxQ1 was confirmed as done in our previous study (10). Cells (5×10^5^ cells/6-cm dish) were harvested by trypsinization and used for RNA isolation and RNA-Seq analysis. The RNA-Seq analysis was performed by Novogene (Sacramento, CA). Other details of RNA-Seq analysis were essentially same as described by us previously (46). Analysis was performed with a combination of software programs including STAR, HTseq, Cufflink, and wrapped scripts. Tophat program was used for alignments and DESeq2 was used for analysis of differential gene expressions. RNA-Seq data presented in this study have been submitted to the Gene Expression Omnibus of NCBI (GSE151059).

### Flow cytometry for analysis of cell cycle distribution and mitotic fraction

EV cells and FoxQ1 overexpressing SUM159 (3 × 10^5^ cells/dish) and MCF-7 cells (6 × 10^5^ cells/dish) were plated in 6-cm culture dishes in triplicate. After overnight incubation, the cells were rinsed with PBS and changed to serum-free medium for 16 hours to synchronize cells in G0/G1 phase. The cells were then released into serum-containing complete medium to resume cell cycle. The cells were harvested with trypsinization at specified time points followed by fixation in 70% ethanol overnight at 4°C. Fixed cells were washed with PBS, stained with propidium iodide (50 μg/mL), and RNase A (80 μg/mL) for 30 minutes and then analyzed using a BD Accuri^™^ C6 flow cytometer (BD Biosciences). For quantitation of the mitotic fraction, cells were plated (SUM159 and HMLE cells - 3 × 10^5^ cells/6-cm dish and MCF-7 cells - 6 × 10^5^ cells/6-cm dish), incubated overnight, and then replaced with fresh medium and incubated for 24 hours. The cells were collected by trypsinization and fixed in 70% ethanol at 4 °C for 2 hours. Subsequently, the cells were permeabilized with 0.25% Triton X-100 for 15 minutes, and incubated with Alexa Fluor 488-conjugated phospho-(Ser10) histone H3 antibody for 1 hour followed by staining with propidium iodide (50 μg/mL) and RNase A (80 μg/mL) for 30 minutes at room temperature. Stained cells were analyzed using BD Accuri™ C6 flow cytometer.

### Western blotting

Details of western blotting have been described by us previously (47). The blots were stripped and re-probed with β-Actin antibody for normalization. Immunoreactive bands were detected by the enhanced chemiluminescence method. Densitometric quantitation was done using UN-SCAN-IT software (Silk Scientific, Orem, UT).

### Quantitative real-time polymerase chain reaction (qRT-PCR)

qRT-PCR was performed as described by us previously (46) and relative gene expression was calculated using the method of Livak and Schmittgen (48). PCR was performed using 2× SYBR green qPCR kit (Thermo Fisher Scientific) with 95°C (15 sec), 60°C (30 sec), and 72°C (20 sec) for 40 cycles. Primers for *IL-8* and *IL-1α* were purchased from GeneCopoeia (Rockville, MD). *VEGFA* was amplified (95°C-15 sec, 60°C-1 min for 40 cycles) with the following primers and *Glyceraldehyde 3-phosphate dehydrogenase (GAPDH*) was used as a normalization control. The primers used were as follows: for *VEGFA* Forward:5’-ATCTTCAAGCCATCCTGTGTG-3’; Reverse; 5’-CAAGGCCCACAGGGATTTTC-3’ and for *GAPDH* Forward: 5’-GGACCTGACCTGCCGTCTAGAA-3’; Reverse; 5’-GGTGTCGCTGTTGAAGTCAGAG-3’.

### ChIP assay

The ChIP assay was performed according to the manufacturers’ protocol (Magnetic Chip kit, Pierce, Rockford, IL) using normal mouse IgG and FoxQ1 antibodies. Other details of ChIP assay have been described by us previously (10). Putative FoxQ1 binding sites at the *IL-8*, *IL-1α*, and *VEGFA* promoters were amplified (60°C, 1 min, 40 cycles) with the following region-specific primers: for *IL-8* site #1, 5-ATGCACTGTGTTCCGTATGC-3’ (forward) and 5’-GCTTTGCTAGTACAGGACAGG-3’ (reverse); site #3, 5’-TGCTTTCTTCTTCTGATAGACCA-3’ (forward) and 5’-TGTTAACAGAGTGAAGGGGCA-3’ (reverse); for *IL-1α*, 5’-TTCTTTGGTGAACTGAGGCA-3’ (forward) and 5’-GTGAGCTGCTATGGAGATGC-3’ (reverse); for *VEGFA* site #1, 5’-AAGGTGAGGCCCTCCAAG-3’ (forward) and 5’-ACCTAGCAGATTGGGGGAAG-3’(reverse); site #2, 5’-GGAGGACAGTTGGCTTATGG-3’(forward) and 5’-GCCAACAGACCTGAAAGAGC-3’(reverse); site #3, 5’-TCCAGATGGCACATTGTCAG-3’(forward) and 5’-TCTGGCTAAAGAGGGAATGG-3’(reverse). Fold enrichment was normalized to the input.

### Analysis of breast cancer TCGA data

RNA-Seq data from TCGA database for breast cancer (n = 1,097) was analyzed using the University of California Santa Cruz Xena Browser (https://xena.ucsc.edu/public/). The correlation coefficient and statistical significance was determined by Pearson test.

### Measurement of IL-8, IL-1α, and VEGFA levels

Cells were plated in 10-cm dishes at a density of 6× 10^5^ (SUM159) or 1 × 10^6^ (MCF-7) cells per dish. The secretion of IL-1α, IL-8, and VEGFA in medium was determined using kits from Quantikine R & D System ELISA kits (Minneapolis, MN) according to the instructions provided by the supplier. The value of IL-1α in EV cells was below detection limit, and therefore a value of 1 was assigned for comparison with FoxQ1 overexpressing SUM159 cells.

### Statistical analysis

Statistical tests were performed using GraphPad Prism (version 7.02). Student’s *t* test was performed for statistical comparisons between two groups.

## 3 Abbreviations

CDK: cyclin-dependent kinase
CDC25C: cell division cycle 25C
DACH1: Dachshund homolog 1
EMT: epithelial-mesenchymal transition
ER: estrogen receptor
EV: empty vector transfected control
GO: gene ontology
HER2: human epidermal growth factor receptor 2
IL: interleukin
KEGG: Kyoto Encyclopedia of Genes and Genomes
PBS: phosphate-buffered saline
PR: progesterone receptor
qRT-PCR: quantitative real-time polymerase chain reaction
TCGA: The Cancer Genome Atlas
VEGF: vascular endothelial growth factor

